# Dynorphin counteracts orexin in the paraventricular nucleus of the thalamus: cellular and behavioral evidence

**DOI:** 10.1101/185520

**Authors:** Alessandra Matzeu, Marsida Kallupi, Olivier George, Paul Schweitzer, Rémi Martin-Fardon

**Affiliations:** Department of Neuroscience, The Scripps Research Institute, La Jolla, CA, 92307

## Abstract

The orexin (Orx) system is known to play a critical role in drug addiction and reward-related behaviors. The dynorphin (Dyn) system, conversely, promotes depressive-like behavior and plays a key role in the aversive effects of stress. Orexin and Dyn are co-released and have opposing functions in reward and motivation in the ventral tegmental area (VTA). Earlier studies showed that microinjections of OrxA in the posterior paraventricular nucleus of the thalamus (pPVT) exerted priming-like effects and reinstated cocaine-seeking behavior, suggesting that Orx transmission in the pPVT participates in cocaine-seeking behavior. The present study sought to determine whether Orx and Dyn interact in the pPVT. Using a cellular approach, brain slices were prepared for whole-cell recordings and to study excitatory transmission in pPVT neurons. The superfusion of OrxA increased spontaneous glutamatergic transmission by increasing glutamate release onto pPVT neurons, whereas DynA decreased glutamate release. Furthermore, the augmentation of OrxA-induced glutamate release was reversed by DynA. To corroborate the electrophysiological data, separate groups of male Wistar rats were trained to self-administer cocaine or sweetened condensed milk (SCM). After self-administration training, the rats underwent extinction training and were tested with intra-pPVT administration of OrxA±DynA under extinction conditions. OrxA reinstated cocaine-and SCM-seeking behavior, with a greater effect in cocaine animals. DynA selectively blocked OrxA-induced cocaine seeking *vs*. SCM seeking. The data indicate that DynA in the pPVT prevents OrxA-induced cocaine seeking, perhaps by reversing the OrxA-induced increase in glutamate release, identifying a novel therapeutic target to prevent cocaine relapse.

## INTRODUCTION

Orexin A (OrxA or hypocretin-1) and orexin B (OrxB or hypocretin-2) are hypothalamic neuropeptides (de Lecea *et al*, 1998; Sakurai *et al*, 1998) that regulate feeding, energy metabolism, and arousal (James *et al*, 2016; Li *et al*, 2016). Two Orx receptors (OxRs) have been identified, OxR1 and OxR2 (Ammoun *et al*, 2003; Sakurai *et al*, 1998). OrxA binds OxR1 and OxR2 with similar affinity, whereas OrxB binds OxR2 with higher affinity (Ammoun *et al*, 2003; Sakurai *et al*, 1998). Orexin-expressing neurons are exclusively found in the hypothalamus (de Lecea *et al*, 1998; Sakurai *et al*, 1998) and project throughout the brain (Peyron *et al*, 1998), playing a key role in modulating reward function, particularly drug-directed behavior (Harris *et al*, 2005). Orexin has been reported to enhance the incentive motivational effects of stimuli that are conditioned to drug availability, increase the motivation to seek drugs of abuse, and enhance the reinforcing actions of drugs of abuse (for review, see (Matzeu *et al*, 2014).

The dynorphin/κ opioid receptor (Dyn/KOR) system is widely distributed in the brain (Watson *et al*, 1982) and is implicated in numerous physiological and pathophysiological processes that are related to mood and motivation (Schwarzer, 2009; Wee and Koob, 2010). The Dyn/KOR system has been suggested to be a putative therapeutic target for the treatment of various neuropsychiatric disorders (Schwarzer, 2009). Two forms of Dyn (DynA and DynB) exist, and both forms (especially DynA) have a marked preference for KORs (Chavkin *et al*, 1982). In contrast to Orx, which promotes arousal and is implicated in the rewarding effects of food, sexual behavior, and drugs of abuse (Tsujino and Sakurai, 2013), Dyn promotes depressive-like behavior and plays a key role in mediating the aversive effects of stress (Bruchas *et al*, 2010). The activation of KORs can attenuate the rewarding effects of drugs of abuse (Wee and Koob, 2010).

Despite their opposing effects on motivation, Orx neurons contain Dyn (Chou *et al*, 2001). Evidence suggests that these peptides act in concert. For example, both Orx and Dyn are co-released during electrical stimulation of the hypothalamus (Li and van den Pol, 2006) and play opposing roles during cocaine self-administration, brain stimulation reward, and impulsivity (Muschamp *et al*, 2014). At the cellular level, OrxA and DynA were applied separately and increased and decreased the firing rate of VTA neurons, respectively (Muschamp *et al*, 2014). Importantly, most recorded VTA neurons were dual-responsive to OrxA and DynA, and the co-application of these peptides resulted in no net changes in neuronal firing, strongly suggesting that OrxA and DynA exert balanced and opposing effects on VTA neurons (Muschamp *et al*, 2014).

The PVT plays a significant role in functions that are related to arousal, attention, and awareness (Van der Werf *et al*, 2002). Data from our laboratory have demonstrated a role for the posterior PVT (pPVT) and particularly Orx transmission in the pPVT in cocaine-seeking behavior (Matzeu *et al*, 2017; Matzeu *et al*, 2016; Matzeu *et al*, 2015). Few γ-aminobutyric acid-ergic neurons are found in local PVT circuitry, indicating that the synaptic network is mostly glutamatergic in this midline nucleus (Arcelli *et al*, 1997). Neurons in the PVT receive one of the densest Orx projections in the brain (Kirouac *et al*, 2005; Peyron *et al*, 1998), and KOR immunoreactivity is high in these neurons (Mansour *et al*, 1996). In the PVT, Orx and Dyn modulate ionic conductance and neuronal activity in opposite directions (Chen *et al*, 2015; Kolaj *et al*, 2014), but the regulation of excitatory synaptic transmission by Orx is poorly understood, and the effect of Dyn is unknown. To determine whether functional cellular interactions exist between Orx and Dyn in the pPVT, whole-cell recordings were performed, and drug-seeking behavior was evaluated.

## MATERIALS AND METHODS

### Rats

Male Wistar rats (*n* = 110; Charles River, Wilmington, MA, USA), weighing 200-225 g (2 months old) upon arrival, were housed two per cage in a temperature-and humidity-controlled vivarium on a reverse 12 h/12 h light/dark cycle with *ad libitum* access to food and water. All of the procedures were conducted in strict adherence to the National Institutes of Health *Guide for the Care and Use of Laboratory Animals* and were approved by the Institutional Animal Care and Use Committee of The Scripps Research Institute.

### Drugs

OrxA and DynA (American Peptide; Sunnyvale, CA, USA) were diluted in 0.9% sodium chloride (Hospira, Lake Forest, IL, USA) and injected in a volume of 0.5 μl. For electrophysiology, OrxA, DynA, and picrotoxin (Sigma, St. Louis, MO, USA) were dissolved in the superfusate. Cocaine (National Institute on Drug Abuse, Bethesda, MD, USA) was dissolved in 0.9% sodium chloride (Hospira, Lake Forest, IL, USA) and administered at a dose of 0.25 mg/0.1 ml. Sweetened condensed milk (SCM; Nestlé, Solon, OH, USA) was diluted in 2:1 (v/v) in water and delivered at 0.1 ml.

### Electrophysiology

pPVT slices were prepared from naive rats (*n* = 10) that were age-matched to the rats that were used in the behavioral experiments. The rats were deeply anesthetized with isoflurane, and brains were rapidly removed and placed in oxygenated (95% O_2_, 5% CO_2_) ice-cold cutting solution that contained 206 mM sucrose, 2.5 mM KCl, 1.2 mM NaH_2_PO_4_, 7 mM MgCl_2_, 0.5 mM CaCl_2_, 26 mM NaHCO_3_, 5 mM glucose, and 5 mM HEPES. Transverse slices (300 μm thick) were cut on a Vibratome (Leica VT1000S, Leica Microsystems, Buffalo Grove, IL) and transferred to oxygenated artificial cerebrospinal fluid (aCSF) that contained 130 mM NaCl, 2.5 mM KCl, 1.2 mM NaH_2_PO_4_, 2.0 mM MgSO_4_·7H_2_O, 2.0 mM CaCl_2_, 26 mM NaHCO_3_, and 10 mM glucose. The slices were first incubated for 30 min at 35°C and then kept at room temperature for the remainder of the experiment. Individual slices were transferred to a recording chamber that was mounted on the stage of an upright microscope (Olympus BX50WI). Recordings were obtained from 8-10 pPVT neurons/condition from slices that were continuously perfused with oxygenated aCSF at a rate of 2-3 ml/min. Neurons were visualized with a 60× water immersion objective (Olympus), infrared differential interference contrast optics, and a charge-coupled device camera (EXi Blue from QImaging, Surrey, Canada). Whole-cell recordings were performed using a Multiclamp 700B amplifier (10 kHz sampling rate, 10 kHz low-pass filter) and Digidata 1440A and pClamp 10 software (Molecular Devices, Sunnyvale, CA, USA). Patch pipettes (4-7 MΏ) were pulled from borosilicate glass (Warner Instruments, Hamden, CT, USA) and filled with 70 mM KMeSO_4_, 55 mM KCl, 10 mM NaCl, 2 mM MgCl_2_, 10 mM HEPES, 2 mM Na-ATP, and 0.2 mM Na-GTP. Liquid junction potential corrections were performed offline. Pharmacologically isolated spontaneous excitatory postsynaptic currents (sEPSCs) were recorded in the presence of picrotoxin (20 μM). Experiments with a series resistance > 15 MΩ or > 20% change in series resistance were excluded from the final dataset. The sEPSC frequency, amplitude, and kinetics were analyzed using MiniAnalysis software (Synaptosoft, Fort Lee, NJ, USA), with measures derived from a minimum time interval of 3 min.

### Behavior

<u>*Self-administration and extinction training (Fig. 1A)*</u>. *Cocaine*. Rats (*n* = 50) were surgically prepared with jugular catheters 7-10 days before cocaine self-administration training in daily 6 h (long access) sessions. Every session was initiated by the extension of two retractable levers into the operant chamber (29 cm × 24 cm × 19.5 cm, Med Associates, St. Albans, Vermont, USA). Responses on the right active lever were reinforced on a fixed-ratio 1 (FR1) schedule by intravenous (IV) cocaine (0.25 mg/0.1 ml/infusion), infused over 4 s followed by a 20-s timeout (TO20) period that was signaled by illumination of a cue light above the active lever. Responses on the left inactive lever were recorded but had no scheduled consequences. *Sweetened condensed milk*. Sweetened-condensed milk (*n* = 50) self-administration was established in daily 30 min sessions on a FR1 TO20 s schedule of reinforcement. Sessions were initiated by the extension of both levers into the operant chamber, and responses on the right active lever resulted in the delivery of SCM (0.1 ml) into a drinking receptacle. Responses on the left inactive lever were recorded but had no scheduled consequences. *Cannulation*. After 14 self-administration sessions, both groups were implanted with a guide cannula (23 gauge, 15 mm, Plastics One, Roanoke, VA, USA) that was aimed at the pPVT (anterior/posterior, −3.3 mm; medial/lateral, ±2.72 mm from bregma; dorsal/ventral, −2.96 mm from dura, at a 25° angle; (Paxinos and Watson, 1997) and positioned 3.5 mm above the target injection point (Fig. 1B). After 7 days of recovery, the animals resumed self-administration training for an additional 7 days. *Extinction*. The day following the completion of self-administration training, the rats were placed under extinction (EXT) conditions (extinction criterion: ≤ 10 active lever presses over the last three sessions). All of the extinction sessions lasted 2 h and were identical to the self-administration sessions but without reward (cocaine or SCM) availability. <u>*Intra-pPVT microinjection*</u>. On the last day of extinction training, each rat received a sham injection for habituation to the microinjection (Fig. 1A). Twenty-four hours later, each animal received an intra-pPVT microinjection of 0.5 μg OrxA (the minimal dose that has been shown to induce cocaine-seeking behavior; (Matzeu *et al*, 2015) ± Dyn-A (0, 1, 2, or 4 μg) or DynA alone (4 μg) using a micro-infusion pump (Harvard 22 Syringe Pump, Holliston, MA, USA). The injectors extended 3.5 mm beyond the guide cannulae. Injections were performed at a flow rate of 0.5 μl/min over 1 min. The injector was kept in place for an additional minute to allow diffusion. Following the injections, the rats were returned to their home cage for 2 min and then transferred to the operant chambers and tested under extinction conditions for 2 h.

**Fig. 1.**
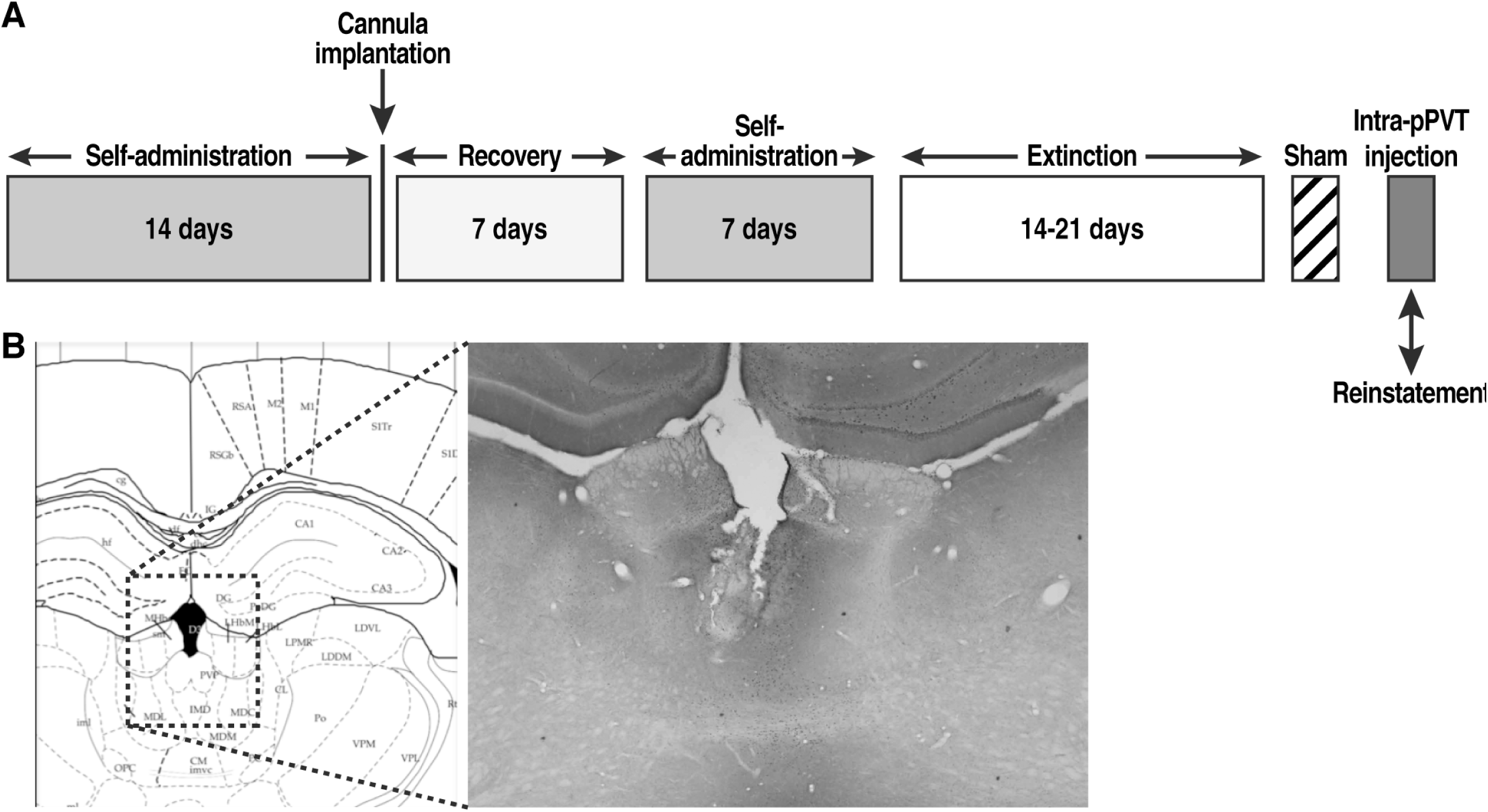
**A**. Behavioral procedure. **B**. Schematic representation of injector placements in the pPVT (Paxinos *et al*, 1997), with photo-representation of the injection track (2.5× magnification).

### Histology

Upon completion of the reinstatement test, the rats were euthanized by CO_2_ inhalation. The brains were then collected and snap frozen. The brains were then sliced into 40 μm coronal sections, and injector placements in the pPVT were verified (Fig. 1B).

### Statistical analysis

The electrophysiology data (mean ± SEM) were analyzed using paired *t*-tests, one-way analysis of variance (ANOVA), or one-way repeated-measures ANOVA followed by the Tukey *post hoc* test when appropriate. For the behavioral data, cocaine and SCM intake during self-administration was analyzed using one-way repeated-measures ANOVA. The number of days to meet the extinction criterion for cocaine and SCM self-administration were compared using unpaired *t*-tests. Differences between extinction, sham, and reinstatement after the vehicle injection were analyzed using two-way repeated-measures ANOVA. Reinstatement following OrxA, DynA, and OrxA+DynA injections was first analyzed by two-way and then individual one-way ANOVAs. Tukey *post hoc* tests were performed to confirm significance.

## RESULTS

Six animals were lost in the cocaine group (two because of health complications, two because of catheter failure, and two because of injection misplacement). One rat was lost in the SCM group (because of injection misplacement), thus reducing the number of animals to 103 (*n* = 10 naive for electrophysiology, *n* = 44 for cocaine, and *n* = 49 for SCM).

### DynA decreased and OrxA increased glutamatergic transmission in naïve pPVT neurons

The modulation of glutamatergic transmission by OrxA and DynA was assessed by studying sEPSCs in pPVT neurons (Fig. 2A). The resting membrane potential was −65 ± 1.4 mV, and the input resistance was 251 ± 19 MΩ. Neurons were held at −68 ± 0.9 mV. DynA (200 nM) significantly decreased the sEPSC frequency of pPVT neurons by 40% ± 6%, from 1.57 ± 0.27 Hz (control) to 0.98 ± 0.22 Hz (paired *t*-test, *t*_9_ = 3.9, *p* < 0.01; Fig. 2B, C). The sEPSC amplitude (34 ± 3 pA), rise time (1.8 ± 0.1 ms), and decay time (3.8 ± 0.5 ms) were not significantly affected by DynA and remained at 94% ± 6%, 97% ± 3%, and 100% ± 8% of control, respectively (Fig. 2D). OrxA (1 μM) significantly increased the sEPSC frequency of pPVT neurons by 38% ± 6%, from 1.49 ± 0.24 Hz (control) to 1.96 ± 0.28 Hz (paired *t*-test, *t*_9_ = 8.7, *p* < 0.001; Fig. 2E). The sEPSC amplitude and rise and decay times were not significantly affected by OrxA and remained at 96% ± 3%, 97% ± 2%, and 96% ± 6% of control, respectively (Fig. 2F).

**Figure 2.**
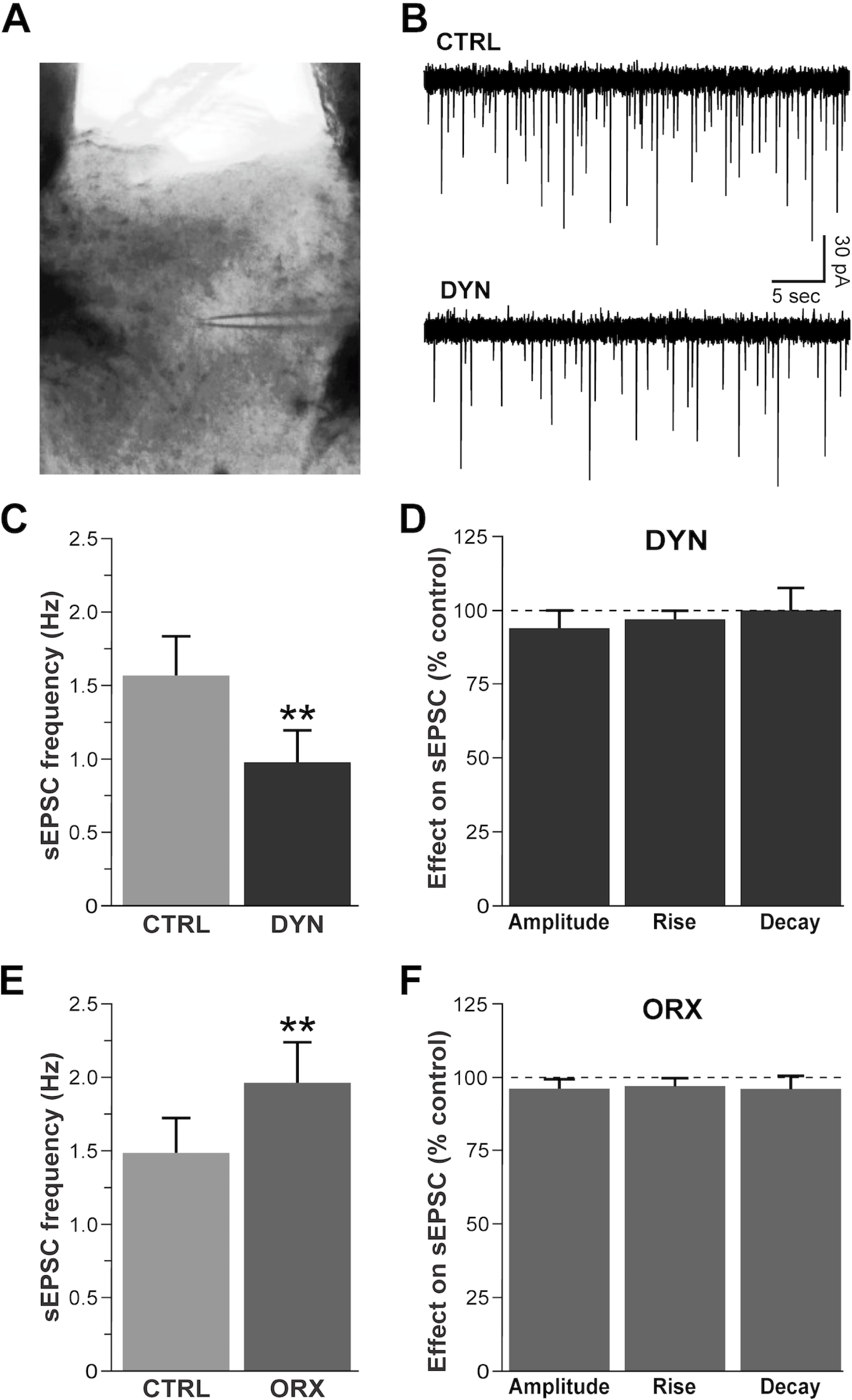
DynA decreased and OrxA increased glutamatergic transmission in pPVT neurons. CTRL, control; DYN, dynorphin-A; ORX, orexin-A. **A**. pPVT slice with the recording electrode (infrared optics, 4× objective). **B**. Representative whole-cell current recordings of sEPSCs (downward deflections). The superfusion of 200 nM DynA decreased the sEPSC frequency by 50%. The resting membrane potential was −64 mV. **C**. DynA significantly decreased the mean sEPSC frequency by 40% in PVT neurons (\*\**p* < 0.01, *vs*. CTRL). **D**. The sEPSC amplitude and rise and decay times were not significantly altered by DynA. **E**. The superfusion of OrxA significantly increased the sEPSC frequency (\*\**p* < 0.01, *vs*. CTRL), an effect that was opposite to DynA. **F**. The sEPSC amplitude and rise and decay times were unchanged by OrxA.

### DynA reversed effects of OrxA on glutamatergic transmission in naïve pPVT neurons

The interaction between OrxA and DynA was studied by first applying OrxA alone and then adding DynA in the continued presence of OrxA. In this neuronal sample, the exposure of pPVT neurons to OrxA (1 μM) significantly increased the sEPSC frequency from 1.58 ± 0.23 Hz to 2.04 ± 0.27 Hz (Tukey *post hoc* test, *p* < 0.01; one-way repeated-measures ANOVA, *F*_2,7_ = 68.1, *p* < 0.01; Fig. 3A, B). The addition of DynA (200 nM) completely reversed the effect of OrxA to control level (sEPSC frequency: 96% ± 8%, control *vs*. OrxA+DynA, *p* > 0.05; Fig. 3). The addition of DynA in the presence of OrxA did not affect the sEPSC amplitude (98% ± 2% of OrxA level), rise time (103% ± 3% of OrxA), or decay time (95% ± 5% of OrxA). The ANOVA of the cumulative data revealed a significant effect of the interaction of the peptides on sEPSC frequency (*F*_2,7_ = 68.1, *p* < 0.01). The Tukey *post hoc* test revealed that the co-application of OrxA+DynA significantly decreased sEPSCs compared with OrxA alone (*p* < 0.01) and significantly increased sEPSCs compared with DynA alone (*p* < 0.01), suggesting that the combination of OrxA+DynA reversed the effect that was observed with each peptide individually (Fig. 3C).

**Figure 3.**
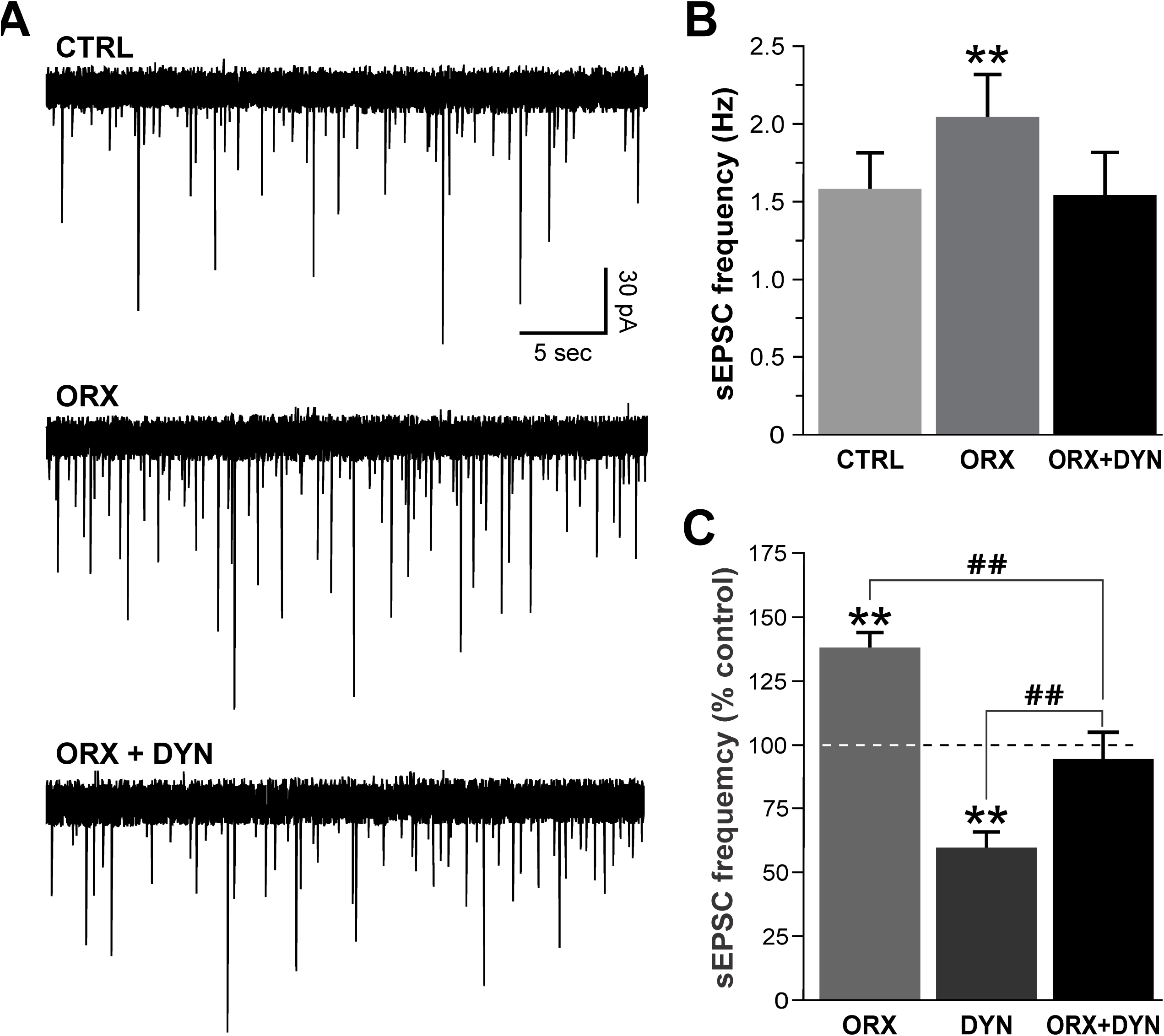
DynA reversed the effect of OrxA on glutamatergic transmission. **A**. The superfusion of 1 μM OrxA increased the sEPSC frequency by 50%, and the further addition of 200 nM DynA in the continued presence of OrxA decreased the sEPSC frequency back to control levels. The resting membrane potential was −69 mV. **B**. OrxA applied alone significantly increased the sEPSC frequency, and the addition of DynA together with OrxA reversed this effect (\*\**p* < 0.01, *vs*. CTRL). **C**. Comparison of sEPSC frequency observed in each experimental group: OrxA alone (38% ± 6% increase, from Fig. 2E), DynA alone (40% ± 6% decrease, from Fig. 2C), OrxA+DynA (4% ± 8% decrease, from Fig. 3B). \*\**p* < 0.01, *vs*. 100% (CTRL); ^##^*p* < 0.01, *vs*. ORX or DYN).

### Cocaine and SCM self-administration training and extinction

In daily 6 h cocaine self-administration sessions, the amount (mg/kg) of cocaine administered gradually increased (repeated-measures ANOVA, *F*_43,860_ = 25.1, *p* < 0.001) from day 1 (15.2 ± 2.3 mg/kg) to day 7 (51.6 ± 4.9 mg/kg) and then remained unchanged until the end of self-administration training (Fig. 4A). In daily 30 min sessions of SCM self-administration, the amount (g/kg) of SCM ingested gradually increased (repeated-measures ANOVA, *F*_48,960_ = 14.3, *p* < 0.001) from day 1 (1.4 ± 0.2 g/kg) to day 7 (7.6 ± 0.5 g/kg) and then remained unchanged until the end of self-administration training (Fig. 4B). By the end of extinction training (19.2 ± 0.3 days for cocaine and 18.8 ± 0.2 days for SCM; unpaired *t*-test, *t*_91_ = 1.01, *p* > 0.05), the rats reached a 3-day average (± SEM) number of responses of 8.4 ± 1.1 for cocaine and 8.2 ± 1.5 for SCM. No differences were observed in the number of responses during extinction, sham, or vehicle injection in the cocaine group *vs*. SCM group (two-way repeated-measures ANOVA; effect of group [cocaine and SCM], *F*_1,18_ = 3.3, *p* > 0.05; effect of time [EXT, SHAM, VEH], *F*_2,36_ = 0.4, *p* > 0.05; group × time interaction, *F*_2,36_ = 1.3, *p* > 0.05; Fig. 5A). Responses on the inactive lever did not change during these procedures (Fig. 5A).

**Fig. 4.**
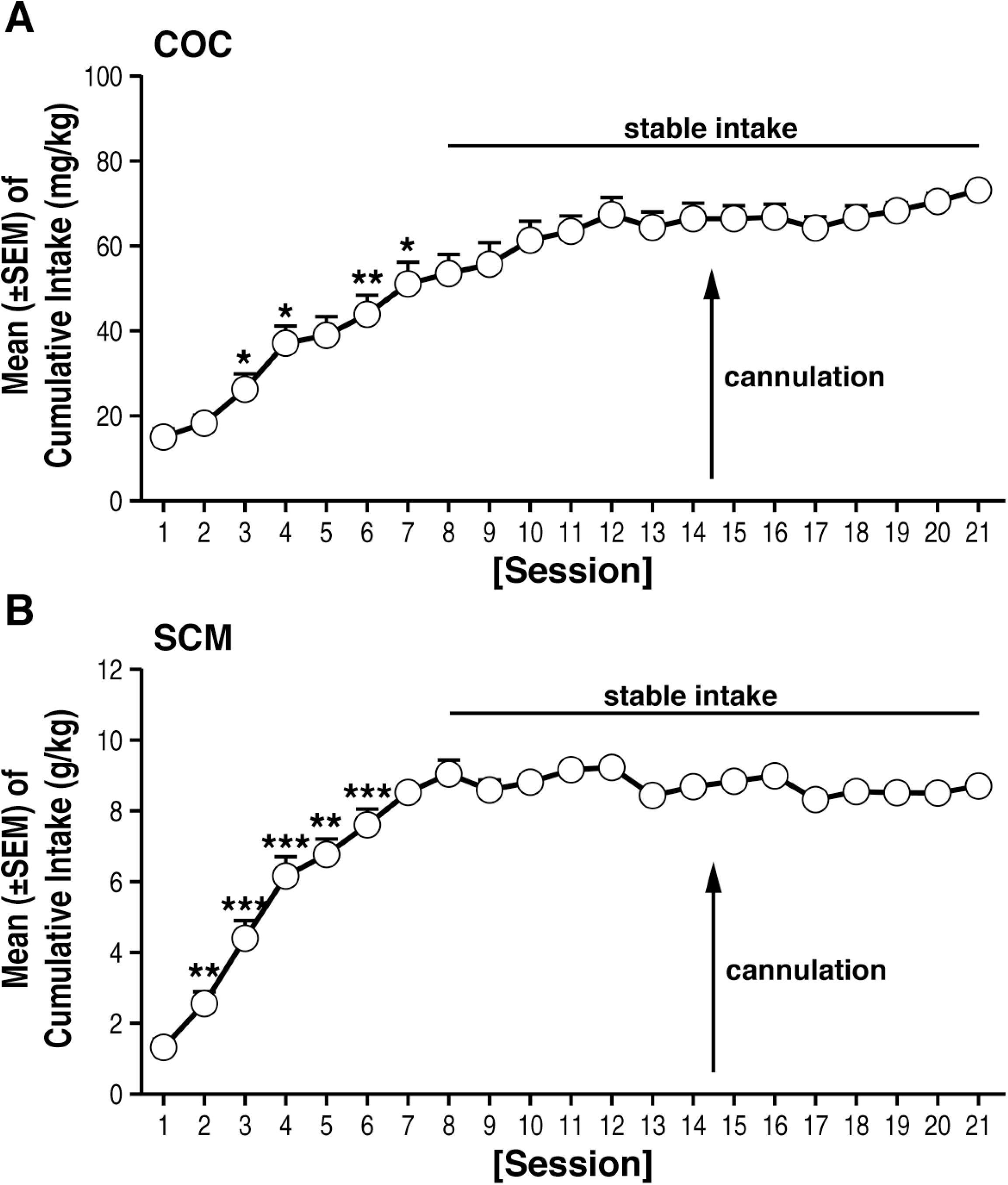
Time course of cocaine (COC) (**A**) and SCM (**B**) self-administration over the 21 days of training. \**p* < 0.05, \*\**p* < 0.01, \*\*\**p* < 0.001, *vs*. respective previous day (Tukey *post hoc* test). The data are expressed as mean ± SEM. *n* = 44 for COC and 49 for SCM.

**Fig. 5.**
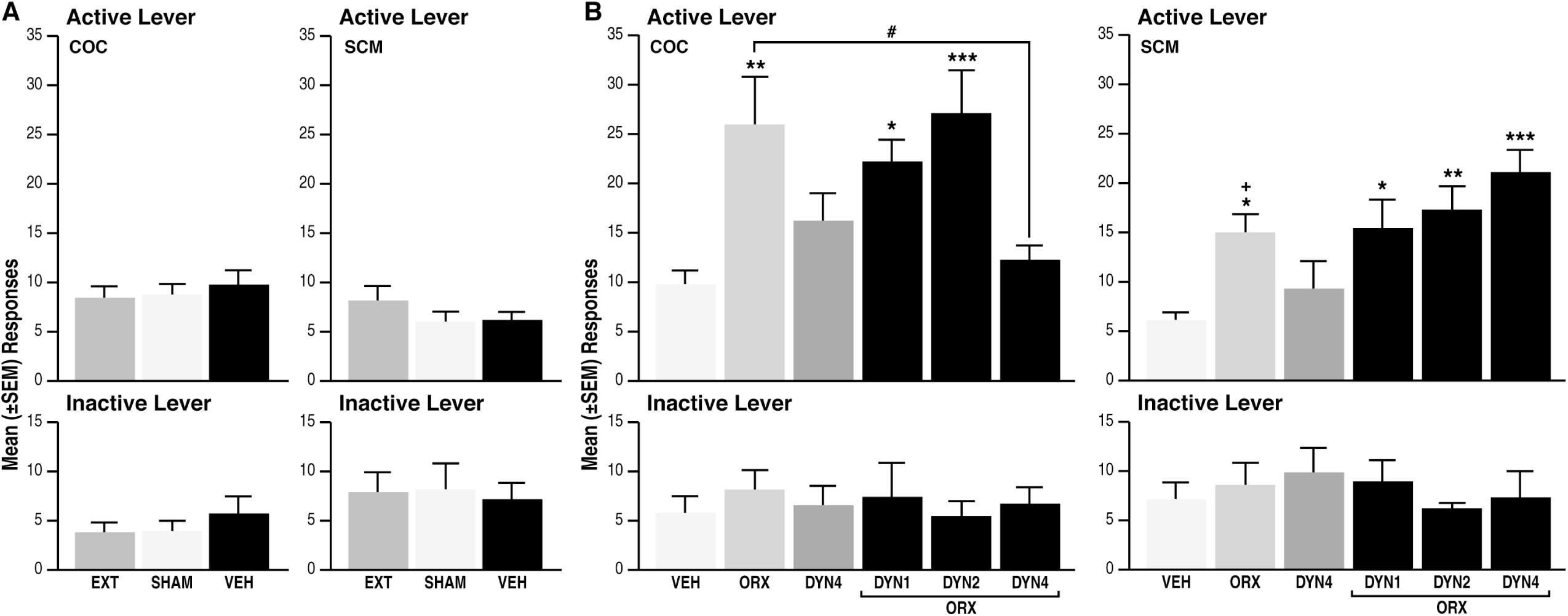
**A**. Cocaine (COC) and SCM animals showed comparable responses during extinction, sham, and vehicle injections. EXT, extinction; SHAM, sham injection; VEH, vehicle; DYN, dynorphin-A; ORX, orexin-A. **B. COC**. Intra-pPVT administration of OrxA but not DynA induced cocaine-seeking behavior. \*\**p* < 0.01, *vs*. VEH. DynA (1 and 2 μg) did not affect OrxA prime-induced reinstatement. \**p* < 0.05, \*\*\**p* < 0.001, *vs*. VEH. DynA (4 μg) reversed OrxA prime-induced reinstatement. ^#^*p* < 0.01, *vs*. ORX. The data are expressed as mean ± SEM. *n* = 9-6 animals/group. **SCM**. Greater OrxA effects in cocaine *vs*. SCM animals. ^+^*p* < 0.05, *vs*. respective treatment in cocaine group. Intra-pPVT administration of OrxA but not DynA induced SCM-seeking behavior. \**p* < 0.05, *vs*. VEH. DynA did not affect OrxA prime-induced reinstatement. \**p* < 0.05, \*\**p* < 0.01, \*\*\**p* < 0.001, *vs*. VEH. The data are expressed as mean ± SEM. *n* = 8-11 animals/group.

### Intra-pPVT administration of OrxA induced cocaine-and SCM-seeking behavior

Following vehicle injections, the mean (± SEM) number of responses was negligible in the two groups (9.8 ± 1.4 for cocaine and 6.2 ± 0.9 for SCM; Fig. 5B). OrxA reinstated (primed) cocaine-and SCM-seeking behavior, with a greater effect in the cocaine group (Tukey *post hoc* test following ANOVA; effect of group [cocaine and SCM], *F*_1,81_ = 10.11, *p* < 0.01; effect of treatment [VEH, ORX 0.5, DYN 4, ORX 0.5+DYN 1, ORX 0.5+DYN 2, and ORX 0.5+DYN 4], *F*_5,81_ = 9.3, *p* < 0.001; treatment × group interaction, *F*_5,81_ = 3.3, *p* < 0.01; Fig. 5B). Responses on the inactive lever remained low and unaltered for both groups at all times (Fig. 5B).

### Intra-pPVT administration of DynA selectively blocked OrxA-induced cocaine-seeking behavior

*Cocaine*. Separate one-way ANOVAs showed that compared with vehicle (n = 9), co-administration of DynA and OrxA (*n* = 7/dose) prevented OrxA-induced cocaine-seeking behavior at the highest dose (4 μg) tested (Tukey *post hoc* test, *p* < 0.05, *vs*. OrxA; ANOVA, *F*_5,38_ = 5.8, *p* < 0.001; Fig. 5B). When administered alone, DynA (*n* = 7) did not produce any effect (*p* > 0.05; Fig. 5B). Inactive lever responses were unaffected and identical across treatments (Fig. 5B). *SCM*. In contrast to the cocaine group, co-administration of increasing doses of DynA (*n* = 6-8 animals/dose) with OrxA did not prevent OrxA’s priming effects compared with vehicle, and OrxA-induced SCM-seeking behavior remained unaffected by DynA (Tukey *post hoc* test, *p* < 0.05, *vs*. vehicle; ANOVA, *F*_5,43_ = 6.5, *p* < 0.001; Fig. 5B). Inactive lever responses were unaffected (Fig. 5B).

## DISCUSSION

The present study provided evidence of a functional interaction between Orx and Dyn in the pPVT at both the cellular and behavioral levels. The data demonstrated that OrxA increased and DynA decreased glutamatergic transmission in the pPVT. When applied together, the peptides counteracted each other’s effects on synaptic activity. Behaviorally, the findings demonstrated that OrxA microinjections in the pPVT reinstated both cocaine-and SCM-seeking behavior, with a greater effect on cocaine-seeking behavior. Moreover, this effect was prevented by DynA only in animals that had a history of extended access to cocaine. As suggested by others (Muschamp *et al*, 2014), the results support the hypothesis that targeting KORs may have beneficial effects for the treatment of disorders that are associated with increased Orx activity.

Electrophysiological investigations have shown that Orx enhances neuronal excitability via pre-and postsynaptic mechanisms (Kukkonen, 2013; Leonard and Kukkonen, 2014). Consistent with the dense projections of Orx neurons to the PVT and mostly glutamatergic nature of the PVT network, we observed an increase in glutamate release by Orx in pPVT neurons that was comparable to the effects that have been reported in the hypothalamus (Li *et al*, 2002; van den Pol *et al*, 1998), laterodorsal tegmental area (Burlet *et al*, 2002), nucleus tractus solitarius (Smith *et al*, 2002), and neocortex (Aracri *et al*, 2015). Previous studies that evaluated the PVT, however, found no effect of Orx on the sEPSC frequency in transgenic mice (Huang *et al*, 2006) or evoked excitatory transmission in rats (Kolaj *et al*, 2007). The use of a different species (Huang *et al*, 2006) or experimental designs (Kolaj *et al*, 2007) may account for such discrepancies. The decrease in glutamatergic transmission that we observed upon DynA application was similar to effects that have been reported in other brain regions (Crowley *et al*, 2016; Wagner *et al*, 1993). Our data showed that OrxA and DynA directly modulated glutamate signaling in an opposite direction, and DynA completely reversed the effect of OrxA, suggesting that these two systems balance each other to regulate neuronal activity in the PVT. Orx and Dyn also have opposing actions on neuronal firing rate in the VTA and counteract each other upon co-release (Muschamp *et al*, 2014). Similarly, Dyn has been shown to inhibit the increase in firing rate that is elicited by Orx in hypothalamic neurons (Li *et al*, 2006) and counterbalance the response to Orx in basal forebrain neurons, preventing overexcitation (Ferrari *et al*, 2016). Importantly, our cellular findings are consistent with the behavioral data, indicating a functional interaction between the Orx and Dyn systems.

The observation that local injections of OrxA in the pPVT induced cocaine-and SCM-seeking behavior strongly supports the hypothesis that Orx projections to the pPVT are important in the modulation of reward-seeking behavior in general. An explanation for the general priming effects of OrxA for both cocaine and SCM could involve the known involvement of the Orx system in arousal. Orexin is well known to regulate general arousal (Sakurai *et al*, 2010), and the anticipation of food reward activate Orx-containing neurons in the PVT (Choi *et al*, 2010). The medial prefrontal cortex is one of the targets of Orx-activated PVT neurons (Huang *et al*, 2006; Ishibashi *et al*, 2005). The present results suggest that Orx inputs to the PVT facilitate cortical activation that is linked to general arousal (Sato-Suzuki *et al*, 2002), which could explain the nonspecific effect of intra-pPVT OrxA injections on both cocaine and SCM reinstatement.

OrxA produced greater reinstatement in the cocaine group *vs*. SCM group, and DynA prevented OrxA-induced cocaine-seeking behavior but had no effect on SCM seeking when injected alone. One possible explanation for the differential effects of OrxA and DynA on cocaine-and SCM-seeking behavior is that cocaine-induced neuroadaptations amplified the motivational effects of OrxA-pPVT signaling following abstinence. The greater OrxA priming effect on cocaine-seeking behavior may also partially explain the greater efficacy of DynA in the cocaine group compared with the SCM group. Orx and Dyn are co-localized and co-released (Chou *et al*, 2001; Li *et al*, 2006), and one may speculate that after long access to cocaine, both the Orx and Dyn systems are sensitized, reflected by a greater priming effect of OrxA and greater inhibitory action of DynA. Dysregulation of the Orx and Dyn systems by cocaine has been reported previously. For example, the Orx system was engaged to a greater extent by drugs of abuse than by SCM in an operant model of reward seeking (i.e., conditioned reinstatement (Martin-Fardon *et al*, 2016; Martin-Fardon and Weiss, 2014; Matzeu *et al*, 2016). These results are consistent with the hypothesis that cocaine produces neuroadaptations in the circuits that control the motivation for natural rewards. Cocaine-induced neuroadaptations of the Dyn system have been well documented. Cocaine enhances Dyn expression, KOR signaling, and KOR levels in the striatum and VTA (Daunais *et al*, 1993; Piras *et al*, 2010; Spangler *et al*, 1996; Unterwald *et al*, 1994) and upregulates *Pdyn* gene expression in the NAC (Caputi *et al*, 2014). κ Opioid receptor downregulation was found in the basolateral amygdala and septum (Bailey *et al*, 2007).

Overall, the present study demonstrates that DynA and OrxA have opposite actions on glutamatergic transmission to balance neuronal activity in the pPVT and control cocaine-seeking behavior. These results suggest that DynA in the pPVT may limit cocaine seeking by at least partially counteracting the OrxA-dependent increase in glutamate release. The present results suggest that manipulating the Dyn/KOR system may be a novel strategy to treat disorders that are associated with increased Orx activity, such as cocaine addiction.

## Funding and Disclosure

This work was supported by the National Institute on Drug Abuse and National Institute on Alcohol Abuse and Alcoholism (grant no. DA033344 and AA024146 to RM-F and AA020608 and AA006420 to OG). The authors declare no conflict of interest.

## Acknowledgements

This is publication number 29472 from The Scripps Research Institute. The authors thank Michael Arends for assistance with manuscript preparation.

